# Corticospinal and corticoreticulospinal projections benefit motor behaviors in chronic stroke

**DOI:** 10.1101/2024.04.04.588112

**Authors:** Myriam Taga, Yoon N. G. Hong, Charalambos C. Charalambous, Sharmila Raju, Leticia Hayes, Jing Lin, Yian Zhang, Yongzhao Shao, Michael Houston, Yingchun Zhang, Pietro Mazzoni, Jinsook Roh, Heidi M. Schambra

## Abstract

After corticospinal tract (CST) stroke, several motor deficits in the upper extremity (UE) emerge, including diminished muscle strength, motor control, and muscle individuation. Both the ipsilesional CST and contralesional corticoreticulospinal tract (CReST) innervate the paretic UE and may have different innervation patterns for the proximal and distal UE segments. These patterns may underpin distinct pathway relationships to separable motor behaviors. In this cross-sectional study of 15 chronic stroke patients and 28 healthy subjects, we examined two key questions: (1) whether segmental motor behaviors differentially relate to ipsilesional CST and contralesional CReST projection strengths, and (2) whether motor behaviors segmentally differ in the paretic UE. We measured strength, motor control, and muscle individuation in a proximal (biceps, BIC) and distal muscle (first dorsal interosseous, FDI) of the paretic UE. We measured the projection strengths of the ipsilesional CST and contralesional CReST to these muscles using transcranial magnetic stimulation (TMS). Stroke subjects had abnormal motor control and muscle individuation despite strength comparable to healthy subjects. In stroke subjects, stronger ipsilesional CST projections were linked to superior motor control in both UE segments, whereas stronger contralesional CReST projections were linked to superior muscle strength and individuation in both UE segments. Notably, both pathways also shared associations with behaviors in the proximal segment. Motor control deficits were segmentally comparable, but muscle individuation was worse for distal motor performance. These results suggest that each pathway has specialized contributions to chronic motor behaviors but also work together, with varying levels of success in supporting chronic deficits.

**Key points summary:**

- Individuals with chronic stroke typically have deficits in strength, motor control, and muscle individuation in their paretic upper extremity (UE). It remains unclear how these altered behaviors relate to descending motor pathways and whether they differ by proximal and distal UE segment.
- In this study, we used transcranial magnetic stimulation (TMS) to examine projection strengths of the ipsilesional corticospinal tract (CST) and contralesional corticoreticulospinal tract (CReST) with respect to quantitated motor behaviors in chronic stroke.
- We found that stronger ipsilesional CST projections were associated with better motor control in both UE segments, whereas stronger contralesional CReST projections were associated with better strength and individuation in both UE segments. In addition, projections of both pathways shared associations with motor behaviors in the proximal UE segment.
- We also found that deficits in strength and motor control were comparable across UE segments, but muscle individuation was worse with controlled movement in the distal UE segment.
- These results suggest that the CST and CReST have specialized contributions to chronic motor behaviors and also work together, although with different degrees of efficacy.

## Introduction

Corticospinal tract (CST) injury from stroke commonly causes a constellation of altered motor behaviors in the upper extremity (UE). These motor deficits include diminished strength, motor control, and muscle individuation (Twitchell, 1951; Brunnstrom, 1966; Lang & Schieber, 2004). In the weeks following stroke, recovery dynamics can diverge not only by deficit but also by UE location, i.e., in the proximal (arm) or distal (hand) UE segment. For example, recovery of motor control plateaus before recovery of strength in the arm (Cortes *et al*., 2017), whereas recovery of both behaviors plateau around the same time in the hand (Cortes *et al*., 2017; Xu *et al*., 2017). The recovery of deficits may also depend on different pathways based on the UE segment. For example, if CST connectivity is disrupted in humans, strength recovery is partial in the hand but robust in the arm (Schambra *et al*., 2019), in keeping with behavioral observations in monkeys after pyramidotomy (Lawrence & Kuypers, 1968; Zaaimi *et al*., 2012; Zaaimi *et al*., 2018). The varying recovery patterns, in term of both deficit and UE segment, suggest the involvement of multiple pathways (Kuypers, 1964; Baker, 2011; Schambra *et al*., 2019).

Two major descending pathways—the injured CST in the ipsilesional hemisphere and the uninjured corticoreticulospinal tract (CReST) in contralesional hemisphere—are potential mediators. Each pathway gains anatomical access to the paretic UE through its descending projections. The injured CST descends mostly contralaterally to the spinal cord, whereas the uninjured CReST descends bilaterally, with an ipsilateral predominance, to the spinal cord (Kuypers, 1981; Lemon, 2008; Fregosi *et al*., 2017). There, both pathways converge onto cervical spinal interneurons and motoneurons that serve the paretic UE (Kuypers, 1960; Kuypers *et al*., 1962; Kuypers, 1981; Riddle *et al*., 2009; Riddle & Baker, 2010). In rodent and monkey models of stroke, both pathways undergo anatomical and functional reorganization following CST injury (Weidner *et al*., 2001; McNeal *et al*., 2010; Zaaimi *et al*., 2012; Bachmann *et al*., 2014; Herbert *et al*., 2015; Darling *et al*., 2018; Ishida *et al*., 2019). The anatomical connectivity and post-stroke changes in these pathways signal their potential roles, not only in early recovery but also in supporting motor behaviors long-term (i.e., the chronic stage of stroke).

By the chronic stage of stroke (i.e. >6 months post-stroke in humans), the recovery of motor behaviors has largely plateaued (Duncan *et al*., 1994; Bernhardt *et al*., 2017). Neurophysiological probes during this time could therefore reveal how pathways support chronic motor behaviors. Transcranial magnetic stimulation (TMS) can be used in chronic stroke to examine ipsilesional CST and contralesional CReST projections to proximal and distal UE muscles (Ziemann *et al*., 1999; McCambridge *et al*., 2020; Taga *et al*., 2021). We recently found that the contralesional CReST had stronger projections to a proximal muscle than a distal one, whereas the lesioned CST had the opposite pattern (Taga *et al*., 2021). These patterns raise the question of their relevance to chronic motor behaviors. Of the chronic stroke studies that examined pathway-behavior relationships, all have focused on only one pathway (Pennisi *et al*., 2002; Thickbroom *et al*., 2002, 2004; Brouwer & Schryburt-Brown, 2006; Cakar *et al*., 2016) or one UE segment (Schwerin *et al*., 2008; Hammerbeck *et al*., 2019). To our knowledge, no chronic stroke studies have systematically examined how each pathway relates to motor deficits in both UE segments.

In addition, it is unclear how effectively chronic motor behaviors are managed in the proximal and distal UE segment. Of the chronic stroke studies that compared motor behaviors between segments, most used subjective clinical scales or unmatched testing paradigms. This has led to conflicting results, with some investigators finding differences in particular segmental behaviors (Lang *et al*., 2006; Cho *et al*., 2012) (Zackowski *et al*., 2004; McPherson & Dewald, 2022) and others finding none (Mercier & Bourbonnais, 2004; Lang *et al*., 2006; Lang & Beebe, 2007). Collectively, these observations call into question whether segmental behavioral differences exist in the chronic phase or are an artifact of the testing instruments.

Identifying pathway relationships to segmental behaviors would advance our understanding of pathway roles in chronic motor function. Moreover, identifying segmental differences in behaviors could indicate the efficacy of pathway support. These insights could guide therapeutic targeting of the pathways in chronic stroke. Therefore, the aims of this study were: (1) to identify how ipsilesional CST and contralesional CReST projections relate to motor behaviors in the proximal and distal segments of the paretic UE, and (2) to identify whether there are segmental differences in motor behaviors in the paretic UE. In a cohort of chronic stroke patients, we tested strength, motor control, and muscle individuation in a representative muscle of the proximal segment (biceps, BIC) and distal segment (first dorsal interosseous, FDI). We used behavioral testing paradigms that were identical across segments to ensure matched samples, and used TMS to characterize CST and CReST projections to each muscle. Healthy controls were studied for normative comparison. We found that each pathway was associated with distinct motor behaviors in both segments, but also cooperated in the proximal segment. Moreover, motor control was segmentally comparable but individuation was worse for the distal segment, indicating that pathway mediation of these behaviors has differing levels of success. These findings suggest that chronic stroke rehabilitation approaches could benefit from targeting each pathway to address specific segmental motor deficits.

## Methods

In this cross-sectional observational study, we examined descending motor neurophysiology and motor behavior in chronic stroke subjects. We studied healthy subjects for normative comparison, with their paretic side designated in a counterbalanced manner. Behavior and neurophysiology were assessed at two visits, 1-2 days apart, with the proximal and distal segment examined at each visit. The orders of the visit and segment testing were randomized. The study was approved by the Institutional Review Board at New York University Grossman School of Medicine (study #18-00959). Subjects were consented in accordance with the Declaration of Helsinki. A subset of the neurophysiological data (stroke N = 8; healthy N = 11) was previously examined (Taga et al., 2021).

### Subjects

All subjects were ≥ 18 years old and were right-handed or ambidextrous (premorbid in stroke). Stroke subjects were included if they had a unilateral ischemic or hemorrhagic supratentorial stroke ≥ 6 months prior resulting in any degree of UE weakness (Medical Research Council score < 5/5 in at least one muscle). Healthy subjects were included if they had a normal motor examination and no history of neurological conditions. Subjects were excluded if they had bihemispheric, cerebellar, or brainstem stroke; traumatic brain injury; musculoskeletal, medical, or non-stroke neurological condition that interferes with motor function; global inattention; legal blindness, visuospatial neglect, or visual field cut greater than a quadrantanopia; seizure history or epilepsy; metal or implanted devices in the head (except mouth) or thorax; participation in another study using investigational therapies or medications; or inability to give informed consent.

We initially enrolled 26 chronic stroke and 35 healthy subjects. Because we sought to compare segmental behavior and neurophysiology from the same individual, we excluded subjects who did not have data from both segments. Reasons for incomplete datasets were inability to perform the task at the correct speed (8 stroke), task performance with significant physiologic tremor (5 healthy, 2 stroke), or persistent muscle fasciculations (1 healthy, 1 stroke). These exclusions ensured that the reported neurophysiology and behavioral findings reflect identical subjects for each segment. The resulting study cohort was 15 mildly-to-moderately impaired stroke subjects and 28 healthy subjects (Table 1).

**Table 1:**
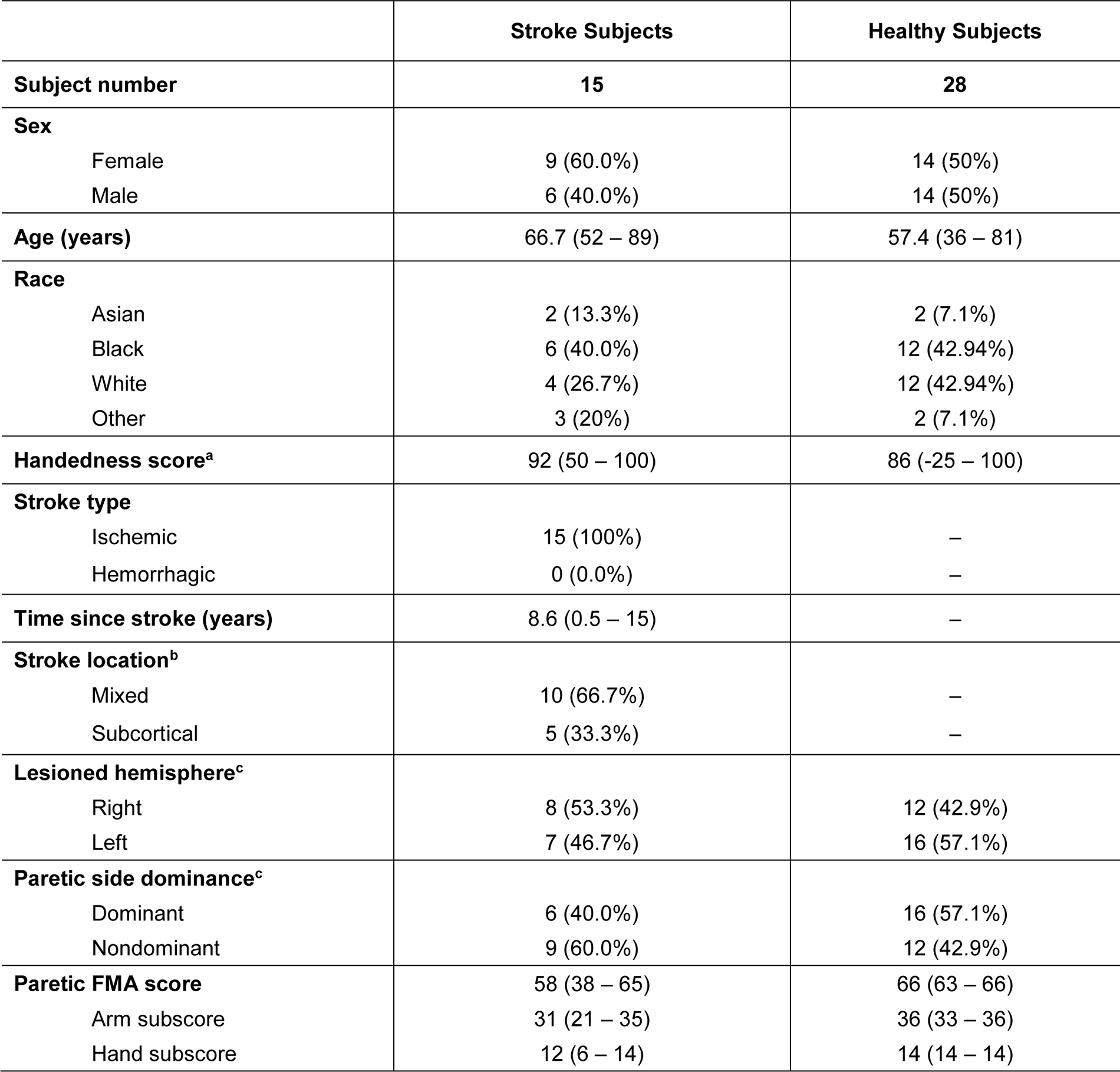
Subject characteristics. Counts (%) or means (ranges) are shown. Stroke subjects were older (p = 0.025) and more impaired (p < 0.0001) than healthy subjects, but were otherwise comparable. ^a^ Edinburgh Handedness Inventory is −100 to +100 with +100 denoting strong right-handedness. ^b^ Stroke location was “mixed” if the lesion involved cortical areas and underlying white matter and “subcortical” if the lesion involved white matter/deep nuclei without cortex. ^c^ Lesioned hemisphere and paretic side were assigned to healthy subjects.

## Experimental procedures

### Motor Behaviors

#### Strength assessment

We used a dynamometer (M550 MyoMeter; Biometrics Ltd, UK) to assess paretic strength (Fig. 1). The dynamometer was fixed to an immobile block or strap and the paretic effector was placed against the dynamometer in a standardized testing position, as described previously (Taga at el., 2021). Subjects performed three trials of maximum elbow flexion or index finger abduction against the dynamometer for 3 s with 30 s rest. We recorded the force (N) generated from the paretic BIC and FDI at 2 kHz with a microprocessor (1401; Cambridge Electronic Design (CED), UK). We used custom scripts offline to extract the mean maximum voluntary force (MVF) force from a 100-ms window centered on the peak force (Signal v7, CED). We computed the average MVF of the three trials.

**Figure 1.**
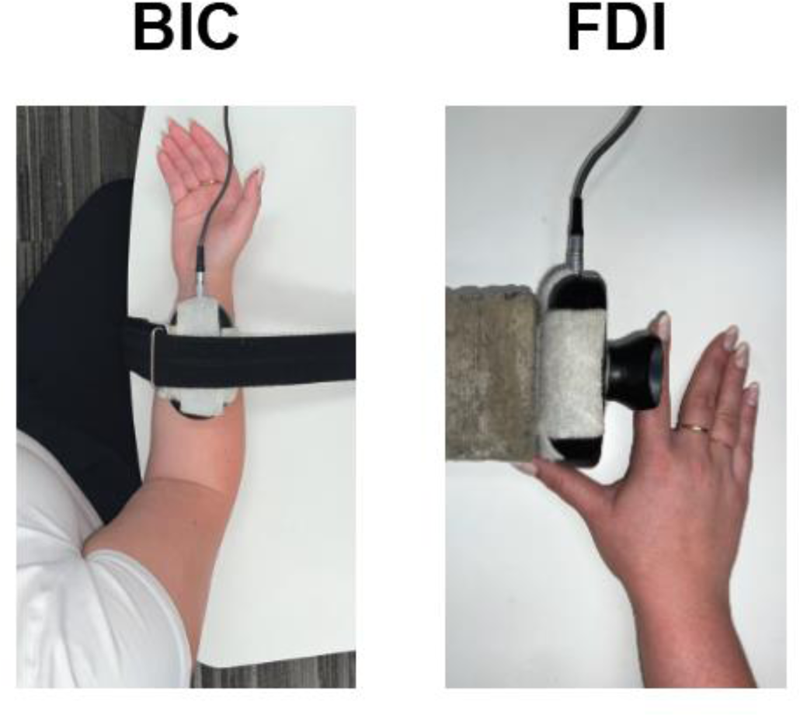
Strength assessment. The experimental set-up for BIC and FDI strength testing is shown. A dynamometer (M550 MyoMeter; Biometrics, UK) was secured to a strap or block, and the paretic effector was placed against it. Subjects were instructed to perform maximal elbow flexions and index abductions in turn against the dynamometer (3 trials; 3 s of contraction; 30 s of rest). The force (N) generated from each muscle was recorded.

#### Motor control assessment

We used a modified motor skill learning task(Shmuelof *et al*., 2012; Shmuelof *et al*., 2014; Gonda *et al*., 2019) to assess paretic motor control (Fig. 2A). Subjects were seated in front of a computer display with their paretic UE positioned on a testing table beside them. Task performance required controlled elbow flexion by the BIC or controlled finger abduction by the FDI. Parts of the UE unnecessary for task performance were braced and restrained against the table. This setup isolated motor control to the instructed muscle and prevented non-instructed muscle activity from intruding on task performance.

**Figure 2.**
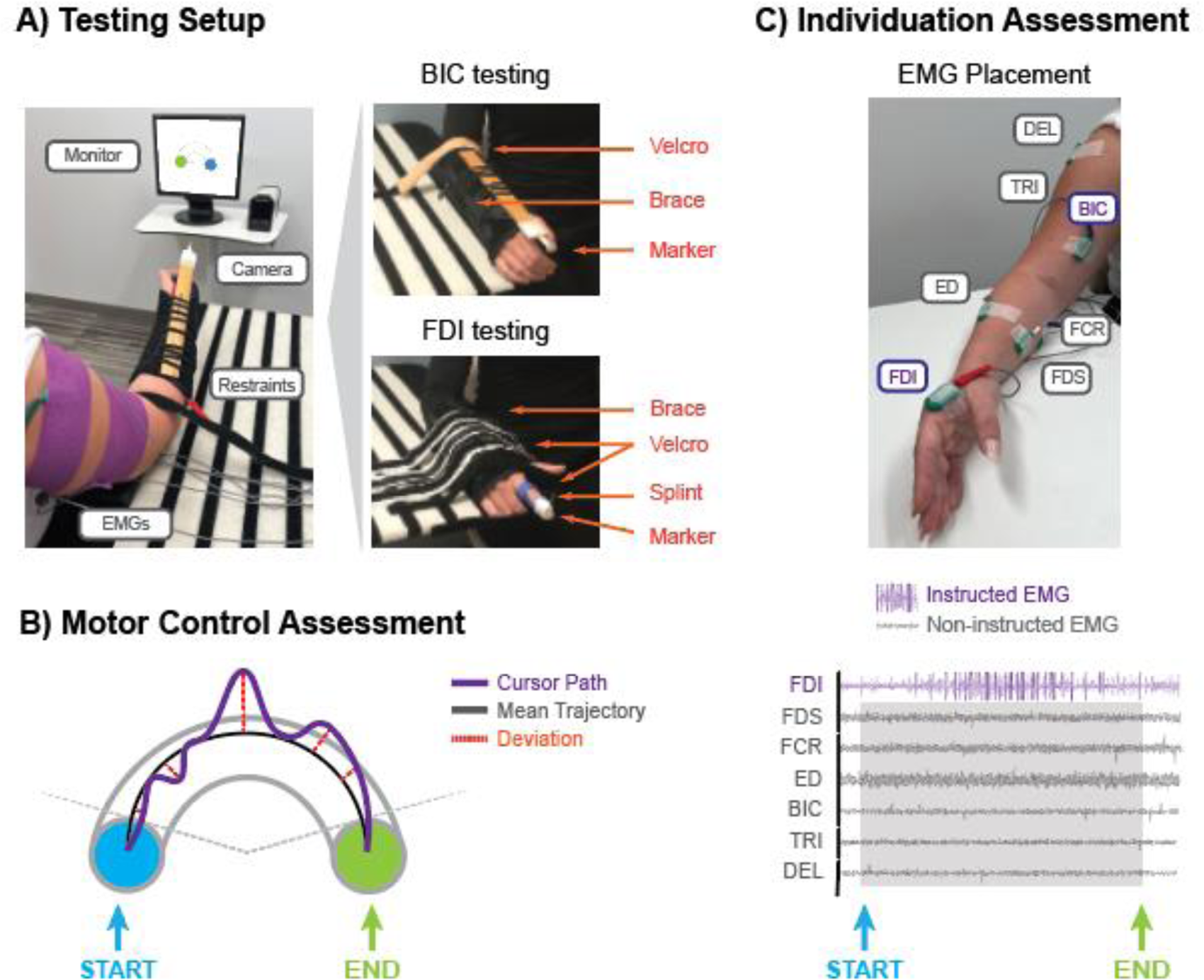
Motor control and individuation assessments. The experimental set-up for BIC and FDI testing is shown. **A) Testing setup.** To assess motor control, subjects were seated in front of a computer screen next to an infrared camera. Surface EMG electrodes were secured to seven UE muscles and the paretic UE was positioned on a testing table. Portions of the UE unnecessary for task performance were restrained using braces and Velcro straps. Movements of the forearm and index finger, respectively actuated by the BIC and FDI, were recorded using reflective marker and the camera. The marker position was shown onscreen as a cursor. **B) Motor control assessment.** Subjects moved the cursor from start (blue) to end (green) of an arc-shaped channel. They were instructed to move as accurately as possible within 1.5 ± 0.3 s. We measured radial oscillations of the cursor path by calculating the mean cursor trajectory (grey line) and the standard deviation (red dotted line) of the distances between the actual cursor path (purple) and this mean trajectory. **C) Muscle individuation assessment.** During arc task performance, we simultaneously recorded EMG from the instructed BIC and FDI and five non-instructed UE muscles. Arc task start and end were demarcated with TTL pulses.

For BIC testing, the paretic UE rested on the table at the elbow and the forearm was elevated off the table. The shoulder was positioned at ∼15° flexion and ∼30° abduction, the elbow at ∼100° flexion, and the forearm midway between neutral and supination. We used a forearm brace (Hely & Weber, US) and a Velcro strap at the elbow to lock out non-instructed wrist and shoulder movement. For FDI testing, the paretic UE rested on the table at the elbow and forearm. The shoulder was positioned as above, with the elbow at ∼70° flexion and the forearm midway between neutral and pronation. We used a finger splint (Rolyan, US), the forearm brace, and Velcro straps over the other fingers, hand, and forearm to lock out non-instructed finger, wrist, elbow, and shoulder movements.

We recorded effector motion with a reflective marker and an infrared camera (Qualisys, Sweden; 100–120 Hz). To ensure identical joint angle excursions across subjects, we equalized effector lengths by adjusting the marker position. For BIC testing, the marker was affixed to a ruler laced into the forearm brace and was adjusted to 45 cm from the lateral epicondyle. For FDI testing, the marker was affixed to a stack of Velcro dots attached to the finger splint and was adjusted to 12 cm from the metacarpophalangeal (MCP) joint. For both setups, the subject was positioned so that the marker was 75 cm from the camera.

Custom software displayed the marker position on a computer monitor along with an arc-shaped channel. The channel (3.7 cm width, 15 cm length) connected a start-circle to an end-circle (each 3.7 cm diameter). The channel was oriented so that effector motion was always toward the subject’s midline, ensuring the same muscles would be activated regardless of left- or right-sided performance. For the arc task, subjects were instructed to move the cursor as accurately as possible, within the channel boundaries, from the start- to the end-circle. They were also instructed to complete the movement within 1.5 ± 0.3 s (average movement speed 120°/s). Prior to each performance block, subjects were shown two computer-generated cursor movements that accurately traversed the arc channel at 120°/s.

Subjects performed two blocks of 15 trials (total of 30 trials) each with ∼1 min rest between. Trial recording began when the cursor was held in the start-circle for 0.2 s, generating a tone and turning the end-circle green. Recording stopped when the cursor was held in the end-circle for 0.2 s, turning it red. Depending on performance, subjects received different feedback at the end of each trial. If the movement speed was in the correct range and the cursor remained inside the arc channel, the cursor path was displayed with a virtual reward (two smiley faces). If movement speed was correct but the cursor left the channel, the cursor path was displayed with red out-of-bound portions and a lesser reward (one smiley face) was shown. If movement speed was incorrect, the cursor path was not displayed and an instruction to “go faster” or “go slower” was shown.

We used custom scripts to inspect recordings and calculate motor performance metrics offline (Igor Pro v8, WaveMetrics Inc, USA). Trials were discarded if the cursor path went off-screen, the camera malfunctioned, or movement speed was incorrect (most common). Analysis was performed on remaining trials (mean count ± SD of included trials: stroke BIC 21 ± 4, stroke FDI 19 ± 7; healthy BIC 23 ± 3, healthy FDI 24 ± 4). The enforced speed range controlled for the speed-accuracy tradeoff (Fitts, 1954). We calculated three motor performance metrics: error rate, path irregularity, and motor control index, as follows.

##### Error rate

We identified trials as accurate or inaccurate based on whether the cursor path remained within the arc boundaries or went out-of-bounds at any point. We computed error rate as the proportion of trials that were inaccurate.

##### Path irregularity

We measured path irregularity—the radial oscillations in the cursor path—to characterize motion smoothness. We did not use jerk (third derivative of position (Flash & Hogan, 1985)) because it is less task-relevant; a path with many accelerations/decelerations can still be successful. Conversely, path irregularity is task-relevant; radial oscillations must be minimized to successfully stay within channel boundaries. To compute path irregularity for each trial, we measured the radial distance from each point on the cursor path to the arc origin and calculated the mean cursor path. We then measured the radial distances between the actual cursor path and mean cursor path, and calculated their standard deviation per trial (Fig. 2B). We normalized the standard deviation to the arc channel width to provide context; for example, a large proportional value implies that radial oscillations were as wide as the arc channel itself. We computed path irregularity as the average of these normalized deviations across trials.

##### Motor control index

We indexed motor control on the arc task using a composite of error rate and path irregularity, because minimizing both parameters is required for successful task execution. For motor control measurement, examining only one parameter may overestimate performance quality; for instance, low error rates can overlook highly irregular cursor paths that remain within the arc channel, whereas low path irregularity can overlook a smooth path that is completely outside of the channel. We accounted for both by computing a motor control index, as follows:

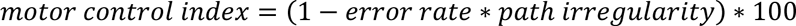

A motor control index nearing 100% indicates superior motor performance, with in-bound and smooth paths. Lower values indicate poorer performance, with out-of-bound and/or wavering paths. For subjects with zero error rate (n; stroke BIC = 1, FDI = 1; healthy BIC = 9, FDI =11), we added a nominal value of 0.01 to avoid zeroing-out path irregularity.

#### Muscle individuation and synergy assessment

We used electromyography (EMG) to assess paretic muscle individuation and synergy expression during performance of the arc task (Fig. 2C). Before we braced and restrained the UE, we placed and secured surface EMG electrodes (SX230-100; Biometrics Ltd., UK) with paper tape and self-adherent wraps. We recorded EMG activity from the ‘instructed’ muscles performing the arc task, the BIC and FDI. We also recorded activity from five ‘non-instructed’ muscles that were not needed for task performance: the anterior middle deltoid (DEL), triceps (TRI), flexor carpi radialis (FCR), extensor digitorum (ED), and flexor digitorum superficialis (FDS). Electrodes (20 mm between poles) were positioned in a muscle belly-tendon orientation using standardized placements (Perotto & Delagi, 2011).

During arc task performance, EMG was amplified x100 (K800; Biometrics Ltd, UK), filtered at 20-460 Hz, and recorded at 2 kHz with a microprocessor and software (Micro 1401-3, Signal v7; CED, UK). TTL pulses were triggered when the cursor in the arc task exited the start-circle and entered the end-circle, marking trial execution on the EMG traces. We used custom scripts (Matlab R2019a) offline to correct DC offsets; to re-filter (high-pass 40 Hz, low-pass 4 Hz) and to rectify the traces; to segment the traces at the TTL marks; and to downsample the traces to 200 equally spaced points per trial. We selected EMG data only from arc task trials performed at the correct movement speed for further analysis. We extracted the maximum EMG amplitude per muscle for each trial. We calculated the average maximum value per muscle during task performance by the BIC or FDI.

Following arc task performance by each effector, we recorded the maximum EMGs that could be elicited in that testing setup, given that joint positioning modifies activity elicitation (Mirka, 1991). Subjects remained in the BIC and FDI testing setups and performed maximum voluntary isometric contractions with each of the seven muscles in turn (3 trials, ∼3 s contraction, ∼60 s rest). EMG was recorded and processed offline as above. We extracted the maximum EMG amplitude in each trial, and calculated the average EMG amplitude across trial for each muscle in each testing setup. We used this maximum EMG amplitude to normalize each muscle’s average EMG amplitudes during arc task performance.

To measure BIC and FDI individuation, we considered how well *non*-instructed muscles *do not* activate when the BIC or FDI is performing the arc task. In addition to the DEL, TRI, FCR, ED, and FDS, we included the FDI as a non-instructed muscle when the BIC performed the task and vice-versa (NB: leaving FDI or BIC out did not change results). We averaged the activity of non-instructed muscles during arc task performance and computed instructed muscle individuation as follows:

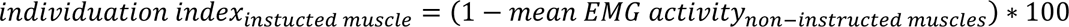

An individuation index closer to 100% indicates that when the instructed muscle performed the task, there was minimal EMG activity in the non-instructed muscles.

##### Non-instructed muscle activity composition

We also explored the composition of non-instructed muscle activity during arc task performance (Fig. 2C). For each subject, we concatenated the normalized EMG data of all trials for each muscle per testing setup. We applied non-negative matrix factorization, which models the EMG patterns as the linear combination of muscle activity vectors multiplied by their temporal activation profile (Lee and Seung 1999). Each activity vector represents the relative level of activation in the muscles, where the magnitudes of activation sum to one. We found that one muscle activity vector explained > 90% of the total data variance based on the global variance accounted for (Roh et al., 2013). From this representation, we extracted a vector representing the activity composition of the six non-instructed muscles. To identify if stroke subjects had abnormal activity compositions, we multiplied their activity vector to each healthy subject vector and averaged the dot products. A mean dot product value closer to 1 indicates that the stroke subject’s non-instructed muscle activity has a composition similar to healthy subjects (Pierella *et al*., 2020). To determine the threshold value that flags abnormal compositions, we multiplied 1000 random six-dimensional unit vectors (MATLAB *rand* function) to generate and rank dot products. Under a dot products threshold of 0.85, stroke subjects have a 95% likelihood that their muscle activity compositions are abnormal.

We further examined whether non-instructed muscle activity during arc task performance was dominated by a flexor or extensor synergy pattern (Twitchell, 1951; Brunnstrom, 1966). We grouped non-instructed muscles by their flexor (BIC, FDS, FCR, DEL) or extensor (FDI, ED, TRI) functions and calculated their mean normalized EMG activity for comparison.

### Neurophysiology

On a separate testing day, we examined ipsilesional CST and contralesional CReST projections using single-pulse TMS, as detailed previously (Taga et al., 2021). Briefly, we used a neuronavigation system (Brainsight; Rogue Research Inc., Canada) and individual structural MRIs (high-resolution 3D T1-weighted images) to place virtual grids over primary and secondary motor areas of each hemisphere (5×5 stimulation sites, 1.25 cm apart, 25 cm^2^) (Fig. 3A). Single-pulse TMS was delivered with a 70 mm figure-of-eight coil and a 200^2^ stimulator (2.2 T output; Magstim Company Ltd., UK). We delivered four TMS stimuli per grid site (100% maximum stimulator output, inter-stimulus interval 7 s) and recorded EMG as above for offline analysis. We used custom Signal scripts to rectify and average the EMG waveforms per grid site, and Matlab scripts to analyze the EMG waveforms.

**Figure 3.**
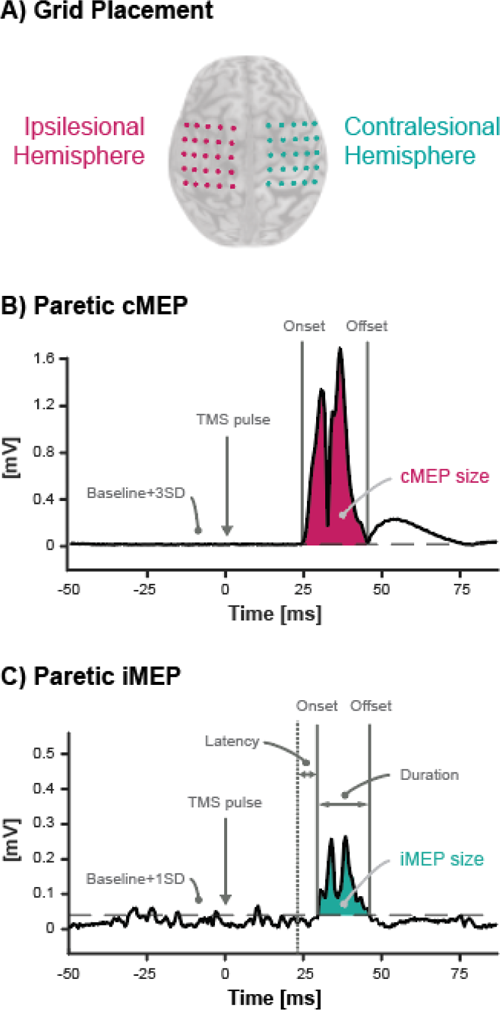
Neurophysiological outcome measures. **A) Grid Placement and assessment.** We placed two virtual grids on the ipsilesional and contralesional hemisphere covering primary and secondary motor regions. To examine injured CST projections, we delivered TMS pulses to the ipsilesional grid while the paretic UE was at rest and recorded contralateral motor evoked potentials (cMEPs). To examine uninjured CReST projections, we delivered TMS pulses to the contralesional grid during paretic muscle activation and recorded ipsilesional motor evoked potentials (iMEPs). **B) Ipsilesional CST projections.** Shown is a cMEP in the paretic FDI of a representative subject. A contralateral waveform qualified as a cMEP when it had a peak amplitude ≥ 0.05 mV, duration ≥ 5 ms, and onset 10-40 ms after TMS. Onset and offset were determined when the waveform crossed 3 standard deviations (SD) of the baseline EMG. **C) Contralesional CReST projections.** Shown is an iMEP in the paretic FDI of a representative subject. An ipsilateral waveform qualified as an iMEP when it had a peak amplitude ≥ 0.05 mV, duration ≥ 5 ms, and an onset 3-10 ms after the reference cMEP (latency delay). Onset and offset were determined when the waveform crossed 1SD of the baseline EMG. The c/iMEP size (mV∗ms) was calculated as the integral of the amplitudes between waveform onset and offset minus the product of the baseline EMG and waveform duration.

To examine the ipsilesional CST, we stimulated the lesioned hemisphere grid while recording from the paretic BIC and FDI; both UEs were at rest (Fig. 3B). We qualified an evoked EMG waveform as a contralateral motor evoked potential (cMEP) if it had a peak amplitude ≥ 0.05 mV, duration ≥ 5 ms, and onset 10-40 ms after TMS. We measured cMEP size (mV∗ms) as the integral of the amplitudes between waveform onset and offset minus the product of the baseline EMG and waveform duration (see (Taga et al., 2021) for additional details). The projection strength of the ipsilesional CST was taken as the maximum cMEP size at the hotspot.

To examine the contralesional CReST, we stimulated the contralesional hemisphere grid while recording from the paretic BIC or FDI during muscle pre-activation; the nonparetic UE was at rest (Fig 3C). Subjects were positioned per strength testing and each muscle was tested independently. Subjects were prompted with verbal and on-screen feedback to generate 20-45% MVF against the dynamometer (∼2 s duration, ∼5 s rest). TMS was delivered during muscle activation, and trials outside of 20-45% MVF range were discarded. Separately, we stimulated the contralesional hemisphere grid while recording from the *non*paretic BIC and FDI; both UEs were at rest. We identified the fastest onset for cMEPs in each muscle, which represented the typical transmission speed for the fast-conducting monosynaptic CST. This reference onset was used to identify and discard fast-onset MEPs that were likely elicited from the ipsilesional hemisphere when a TMS coil wing was near the midline on the contralesional hemisphere. Since the CReST is a slower-conducting oligosynaptic pathway, a true ipsilateral MEP (iMEP) from the contralesional hemisphere should arrive at a delay.

We qualified an evoked waveform as an iMEP if it had a peak amplitude ≥ 0.05 mV, duration ≥ 5 ms, and onset 3-10 ms after the reference cMEP. We measured the iMEP size (mV∗ms) per cMEP measurement and used the largest iMEP to identify the contralesional CReST hotspot for each muscle. At each muscle’s iMEP hotspot, we recorded 20 additional stimuli at 20-45% MVF. If no iMEPs were initially identified (stroke: BIC 8, FDI 1; healthy: BIC 14, FDI 3), we used the cMEP hotspot identified with the nonparetic recordings above. The projection strength of the contralesional CReST was taken as the average iMEP size of all trials with iMEPs at the hotspot.

Muscles with absent cMEP or iMEPs were assigned an MEP size of 0 mV∗ms. We also categorized functional pathway projections to the muscles as present (MEP+) or absent (MEP-).

### Clinical and psychometric assessments

At the behavioral testing visit, we measured clinical motor impairment with the UE Fugl-Meyer Assessment (FMA) (Fugl-Meyer et al., 1975), calculating total UE FMA score (66 points maximum) and subscores for the arm (36 points) and hand (14 points). Following each visit, we used a visual analog scale (VAS) to quantify alertness, excitement, discomfort, fatigue, sleep duration, and alcohol and caffeine consumption in the previous 24 hours.

### Statistical analyses

We first tested for differences in demographic, clinical, and psychometric data between subject groups. We applied the Anderson-Darling test to assess normal distribution, revealing that behavior and neurophysiology data deviated from it. We used independent Student’s t-tests for continuous data, Wilcoxon Rank Sum tests (i.e. Mann Whitney U) for ordinal and non-normally distributed data, and Fisher’s exact tests for nominal data.

Motor behaviors and neurophysiology were non-normally distributed and required non-parametric tests. We first examined behavioral differences between subject groups and UE segments. We compared motor behaviors between stroke and healthy subjects using Wilcoxon Rank Sum tests. We compared motor behaviors between proximal and distal segments using Wilcoxon Signed Rank tests for matched pairs.

We next examined how CST and CReST projection strengths related to motor behaviors in each segment. We correlated cMEP or iMEP size to motor behaviors using Spearman’s rank correlation. We also explored if functional pathway connectivity influenced motor behavior in stroke, splitting muscles by MEP presence/absence and comparing behaviors with Wilcoxon Rank Sum tests.

We performed all analyses in JMP Pro 16 (SAS Institute Inc) and SPSS (IBM SPSS Statistics 28). P-values were uncorrected with significance set at α=0.05. Given data non-normality, group summaries are reported as median and interquartile range unless otherwise stated.

## Results

### Demographic and clinical characteristics

Group data for stroke and healthy subjects are shown in Table 1. Subject groups were comparable demographically (all p > 0.282) except for age, where stroke subjects were older than healthy subjects by 9.3 years on average (t_30_ = 2.4, p = 0.025). As expected, stroke subjects had greater UE impairment than healthy subjects, evidenced by lower total UE FMA scores (z = 5.5, p < 0.0001) and lower FMA subscores in the arm (z = 5.5, p < 0.0001) and hand (z = 4.2, p < 0.0001).

Psychometric data from the assessment visits are shown in Table 2. Stroke subjects were more excited to participate at both visits (behavior, z = 2.2, p = 0.028; TMS, z = 3.3, p = 0.0009) and were more alert during the TMS visit (z = 2.1, p= 0.037) than healthy subjects. Healthy subjects showed a trend for more alcohol consumption than stroke subjects in the 24 h preceding the behavior visit (z = 1.7, p = 0.090). There were no other differences between subject groups in levels of discomfort, fatigue, sleep duration, or caffeine consumption (all z > 1.3, p > 0.210).

**Table 2:**
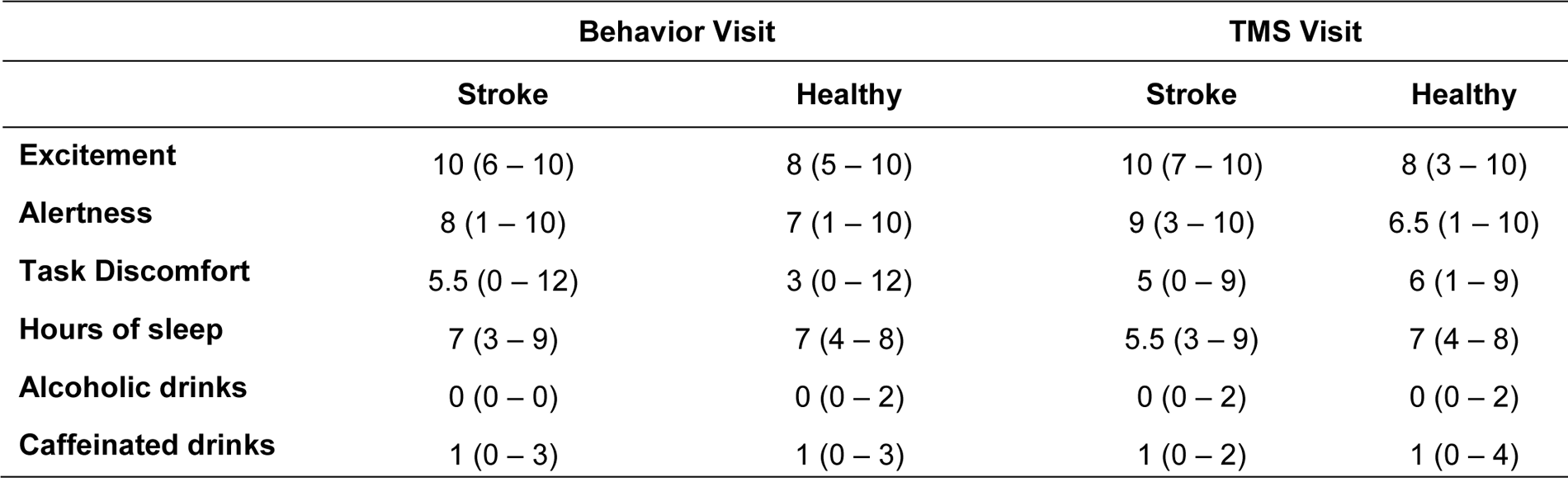
Psychometric measurements at the behavior and TMS testing visits. Median (range) shown. The VAS is from 1–10, with 10 highest. Alcoholic and caffeinated drinks were counted from the 24 hours preceding the visit. Stroke subjects were more excited to participate at both visits (p < 0.028) and were more alert for the TMS visit (p = 0.037).

|  | Behavior Visit |  | TMS Visit |  |
| --- | --- | --- | --- | --- |
|  | Stroke | Healthy | Stroke | Healthy |
| **Excitement** | 10 (6 – 10) | 8 (5 – 10) | 10 (7 – 10) | 8 (3 – 10) |
| **Alertness** | 8 (1 – 10) | 7 (1 – 10) | 9 (3 – 10) | 6.5 (1 – 10) |
| **Task Discomfort** | 5.5 (0 – 12) | 3 (0 – 12) | 5 (0 – 9) | 6 (1 – 9) |
| **Hours of sleep** | 7 (3 – 9) | 7 (4 – 8) | 5.5 (3 – 9) | 7 (4 – 8) |
| **Alcoholic drinks** | 0 (0 – 0) | 0 (0 – 2) | 0 (0 – 2) | 0 (0 – 2) |
| **Caffeinated drinks** | 1 (0 – 3) | 1 (0 – 3) | 1 (0 – 2) | 1 (0 – 4) |

### Motor behavior in stroke versus healthy subjects

We first evaluated if stroke subjects had appreciable motor deficits with respect to healthy subjects. Motor behaviors by subject group and segment are shown in Table 3 and individual subject data are shown in figures.

**Table 3.**
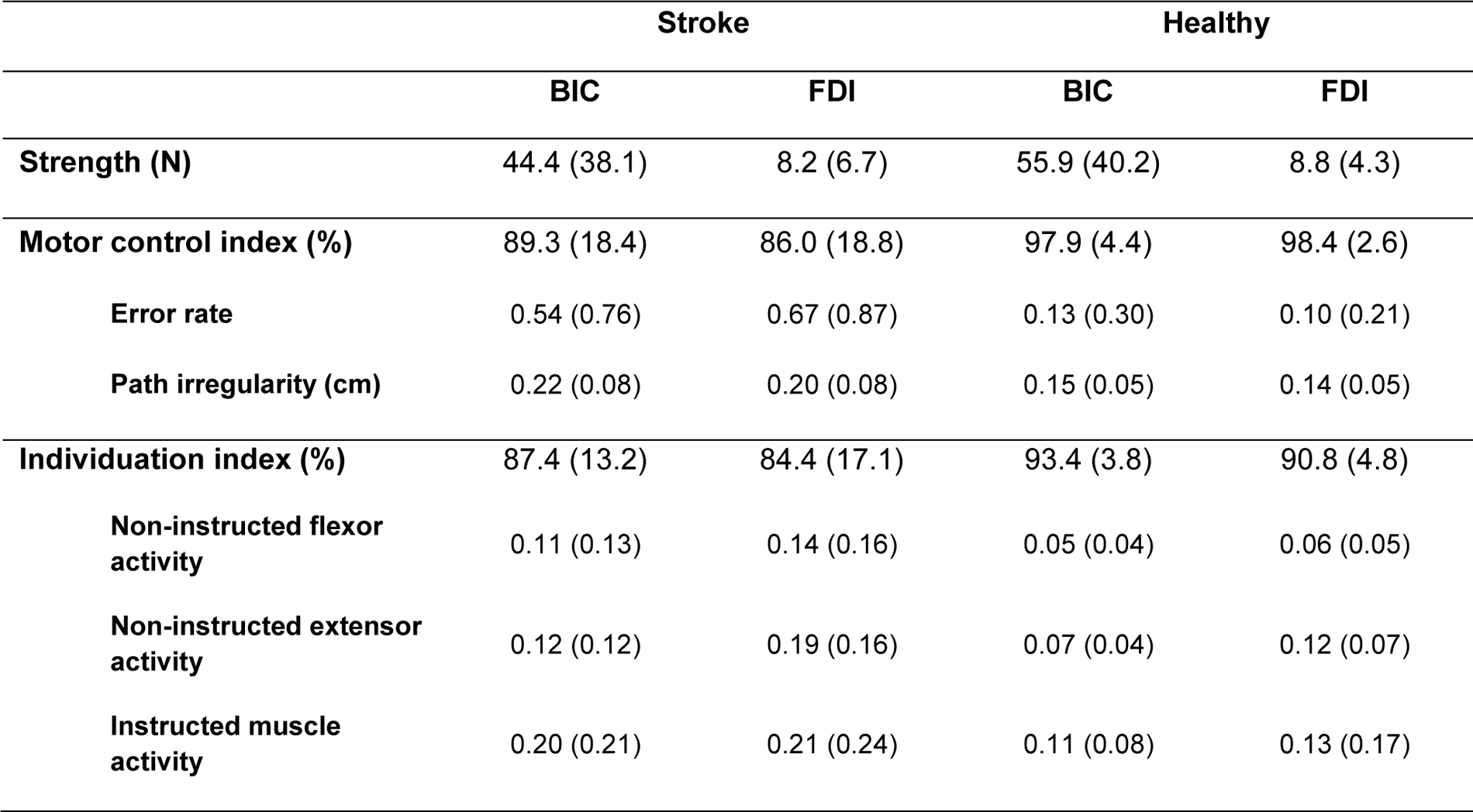
Motor behavior outcomes by subject group and muscle. Median (IQR) shown. The motor control index captures the error rate and path irregularity of task performance, where values closer to 100% indicate accurate, smooth movements. The individuation index captures non-instructed muscle activity during task performance, where values closer to 100% indicate that non-instructed muscles are appropriately inactive. Muscle activity values are normalized to the muscle’s maximum EMG value.

|  | Stroke |  | Healthy |  |
| --- | --- | --- | --- | --- |
|  | BIC | FDI | BIC | FDI |
| **Strength (N)** | 44.4 (38.1) | 8.2 (6.7) | 55.9 (40.2) | 8.8 (4.3) |
| **Motor control index (%)** | 89.3 (18.4) | 86.0 (18.8) | 97.9 (4.4) | 98.4 (2.6) |
| **Error rate** | 0.54 (0.76) | 0.67 (0.87) | 0.13 (0.30) | 0.10 (0.21) |
| **Path irregularity (cm)** | 0.22 (0.08) | 0.20 (0.08) | 0.15 (0.05) | 0.14 (0.05) |
| **Individuation index (%)** | 87.4 (13.2) | 84.4 (17.1) | 93.4 (3.8) | 90.8 (4.8) |
| **Non-instructed flexor activity** | 0.11 (0.13) | 0.14 (0.16) | 0.05 (0.04) | 0.06 (0.05) |
| **Non-instructed extensor activity** | 0.12 (0.12) | 0.19 (0.16) | 0.07 (0.04) | 0.12 (0.07) |
| **Instructed muscle activity** | 0.20 (0.21) | 0.21 (0.24) | 0.11 (0.08) | 0.13 (0.17) |

We first compared segmental strength between subject groups (Fig. 4). Strength was modestly lower in both muscles for stroke subjects, but group-level differences were not significant in either muscle (BIC, z = 0.98, p = 0.327; FDI, z = 0.96, p = 0.339).

**Figure 4.**
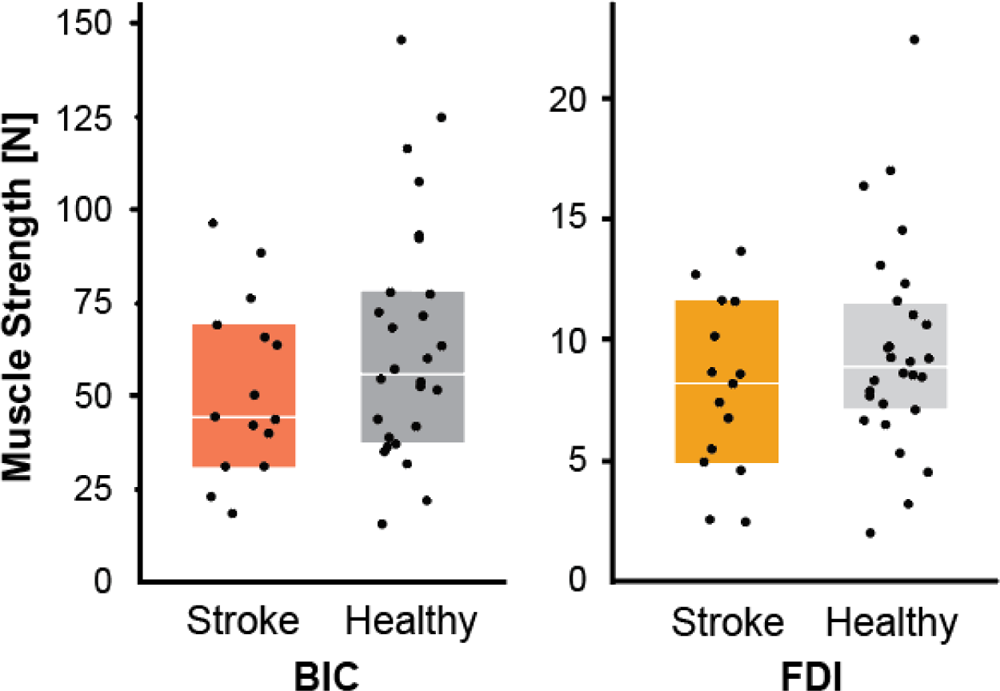
Muscle strength is comparable between stroke and healthy subjects. The box plots represent median values with upper and lower quartiles, and single dots represent single subjects. We assessed differences in BIC and FDI strength between subject groups but found no significant differences in either segment (p > 0.327). Within subject groups, BIC and FDI were not compared directly because of inherent differences in muscle strength (note axes differences).

We then compared segmental motor control between subject groups (Fig. 5). Stroke subjects had worse task performance than healthy subjects in terms of higher error rates (BIC, z = 2.8, p = 0.004; FDI, z = 3.1, p = 0.002) and greater path irregularity (BIC, z = 3.6, p < 0.001; FDI, z = 3.4, p < 0.001). Accordingly, stroke subjects had lower motor control indices than healthy subjects in both muscles (BIC, z = 3.2, p = 0.002; FDI, z = 3.3, p = 0.001).

**Figure 5.**
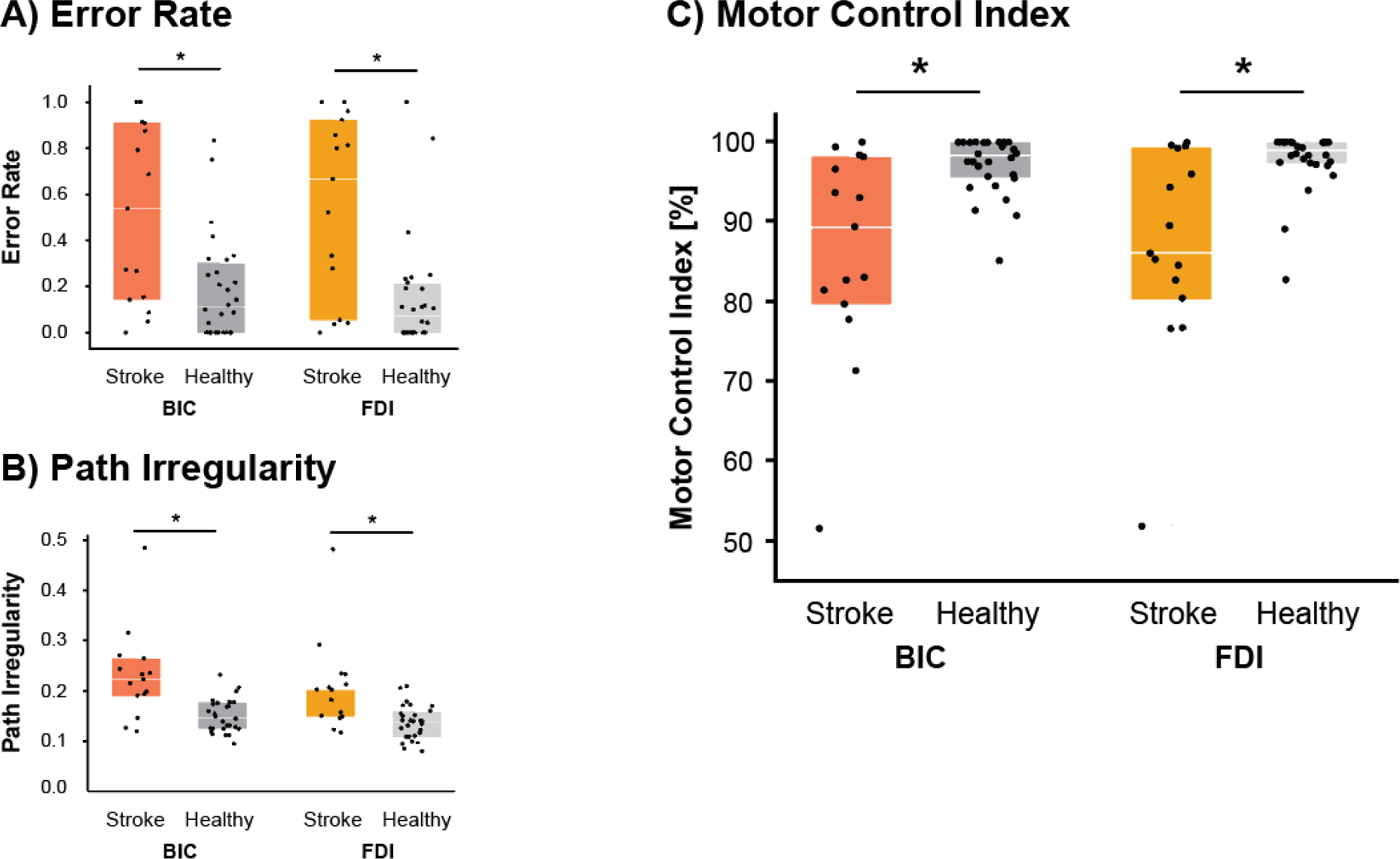
Motor control is worse in stroke subjects, but comparable between UE segments. The box plots represent median values with upper and lower quartiles, and single dots represent single subjects. **A) Error rate.** Stroke subjects had higher error rates than healthy subjects in both UE segments (BIC, p = 0.004; FDI, p = 0.001). **B) Path irregularity.** Stroke subjects had significantly greater path irregularity than healthy subjects in both UE segments (BIC, p = 0.0003; FDI, p = 0.001). **C) Motor control.** Stroke subjects had lower motor control indices than healthy subjects in both segments (BIC, p = 0.002; FDI, p = 0.0007). Within subject groups, motor control was comparable between proximal and distal UE segments (stroke, p = 0.570; healthy, p = 0.084). * p < 0.05

We also compared segmental muscle individuation between subject groups (Fig. 6A). Stroke subjects had worse individuation indices than healthy subjects during task performance by the instructed muscle (BIC, z = 3.2, p = 0.002; FDI, z = 3.7, p < 0.001). Stroke subjects also needed relatively more activation of their instructed muscle than healthy controls to perform the arc task (BIC, z = 2.2, p = 0.026; FDI, z = 2.7, p = 0.007; Fig. 6B).

**Figure 6.**
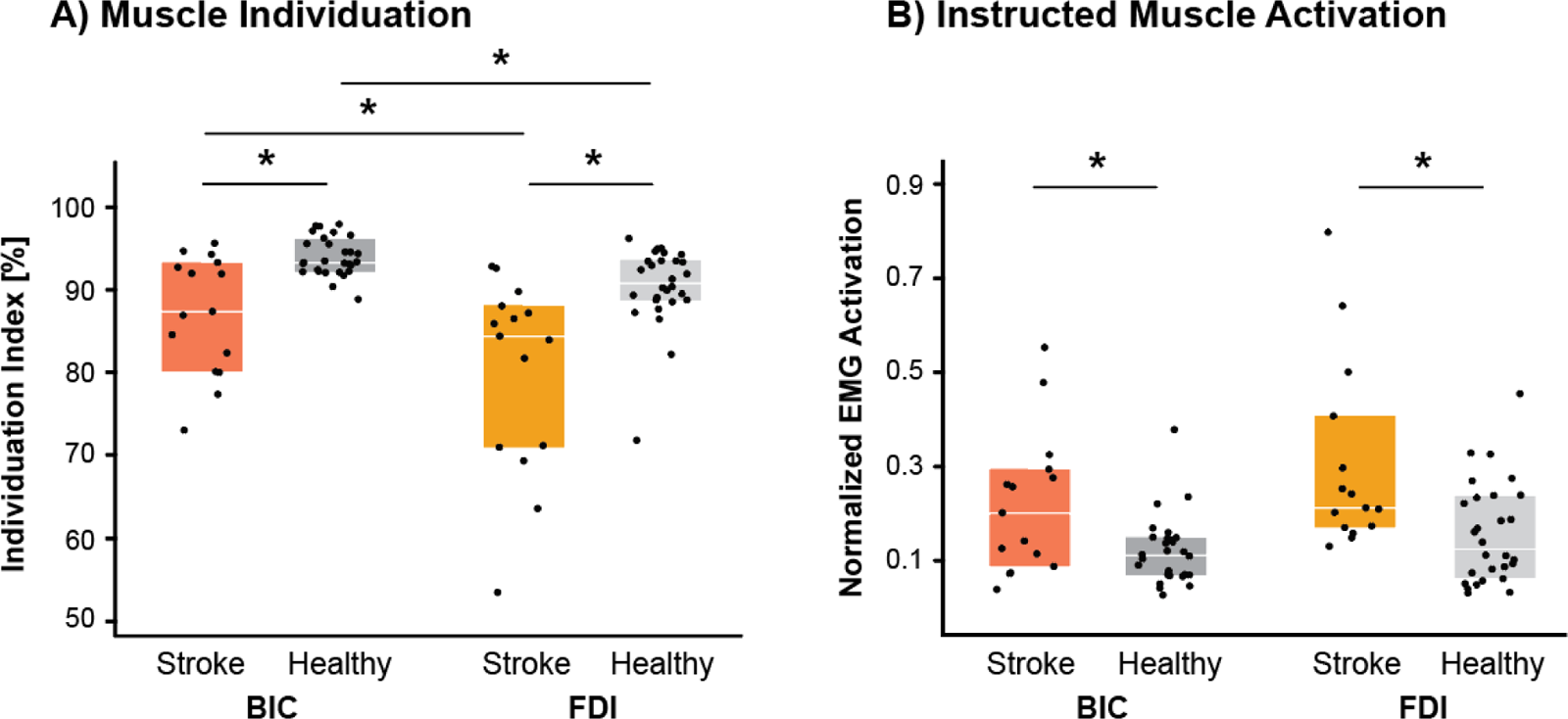
Muscle individuation is worse in stroke subjects and in the distal UE segment. The box plots represent median values with upper and lower quartiles, and single dots represent single subjects. We assessed differences in muscle activation and individuation between subject groups and UE segments. **A) Muscle individuation.** Stroke subjects had lower individuation indices than healthy subjects during task performance by the instructed muscle (BIC, p = 0.002; FDI, p < 0.001). Within subject groups, muscle individuation was worse with task performance by the FDI than the BIC (stroke, p = 0.003; healthy, p < 0.001). **B) Instructed muscle activation.** Stroke subjects needed relatively more activation of their instructed muscle than healthy controls to perform the arc task (BIC, p = 0.026; FDI, p = 0.007). In stroke subjects, there was no significant difference between BIC and FDI activation during task performance (p = 0.256), indicating that the lifting the weight of the effector was similarly demanding for both muscles. In healthy subjects, there was a strong trend for a higher FDI than BIC activation during task performance (p = 0.062). * p < 0.05

Finally, we compared the compositions of non-instructed muscle activation between groups, first examining the dot products of their muscle activity vectors. Mean dot products between stroke and healthy subjects were below 0.85 for each task (BIC 0.61 (0.09), FDI 0.67 (0.06)), indicating abnormal compositions of non-instructed muscle activity. We explored whether these abnormal compositions reflected specific activation of flexor or extensor synergies. We found instead more generalized non-instructed muscle activation: stroke subjects had greater activation during BIC performance of both flexors (z = 3.4, p = 0.0008) and extensors (z = 2.6, p = 0.009). Similarly, stroke subjects had higher activation during FDI performance of flexors (z = 3.2, p = 0.002) and extensors (z = 3.4, p = 0.0006).

Collectively, these results indicate that our stroke cohort had deficits in motor control, muscle individuation, and muscle co-activation patterns in both segments of the paretic UE, despite relatively preserved strength.

### Motor behavior in stroke: proximal versus distal UE segment

In stroke subjects, we evaluated if motor deficits differed by UE segment. We compared motor control, muscle individuation, and non-instructed flexor and extensor activity (strength could not be normalized and was therefore not segmentally compared.) To ensure that segmental comparisons would not be influenced by differential performance demands, we imposed identical task parameters for data acquisition. However, it is conceivable that the mass the effector (i.e. index finger vs. forearm) drove different degrees of muscle activation, which could influence individuation (Xu *et al*., 2017). We confirmed that there was no significant difference in BIC and FDI activation during task performance (z = 1.1, p = 0.256), indicating that the lifting the weight of the effector was similarly demanding for both muscles.

We found that motor control did not differ between the muscles (z = 0.6, p = 0.570; Fig. 5C). However, muscle individuation was worse during task performance by the FDI than BIC (z = 3.0, p = 0.003; Fig. 6A). Moreover, non-instructed flexors and extensors were comparably active during BIC task performance (z = 0.5, p = 0.650), whereas extensors were more active than flexors during FDI task performance (z = 2.8, p = 0.005). These results indicate that motor performance by the FDI provoked stronger, extensor-dominated muscle activity, while BIC performance provoked weaker and more evenly distributed muscle activity.

### Relationships between descending motor pathways and behavior in stroke

Next we examined how the projection strengths of the ipsilesional CST and contralesional CReST related to segmental motor behaviors in the paretic UE. We correlated cMEP and iMEP size with strength, motor control, and individuation in the tested muscle.

#### Muscle strength

In the proximal segment, CST projection strength significantly related to BIC strength, with larger cMEPs associated with higher MVFs (ρ = 0.53, p = 0.045; Fig. 7A). CReST projection strength also significantly related to BIC strength, with larger iMEPs associated with higher MVFs (ρ = 0.84, p < 0.0001; Fig. 7A).

**Figure 7.**
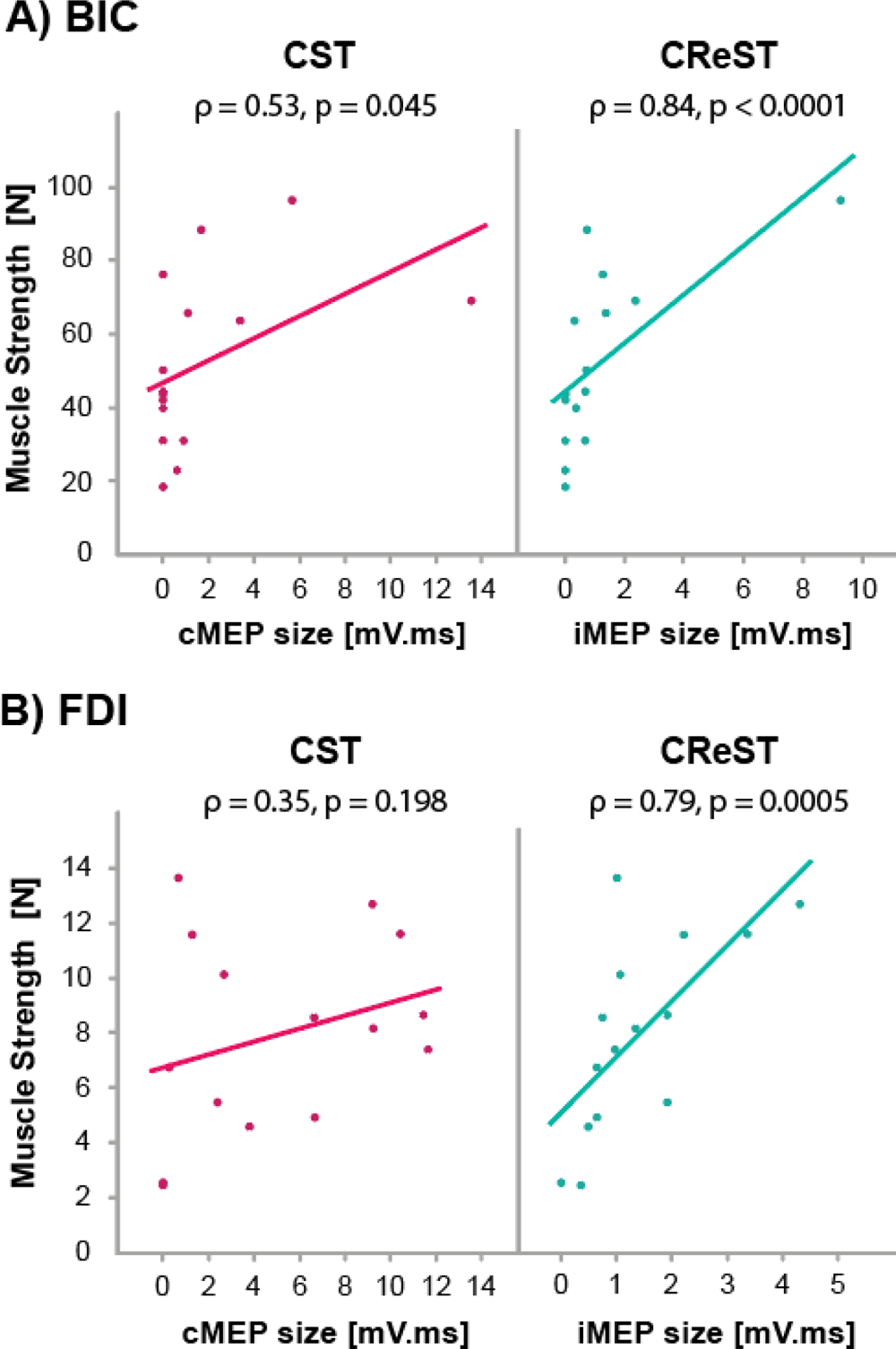
Contralesional CReST projections strongly relate to paretic muscle strength. **A) In BIC**, CST and CReST projection strengths significantly related to muscle strength. Larger cMEPs were associated with higher MVFs (p = 0.045) and larger iMEPs also were associated with higher MVFs (p < 0.0001). Removal of the subject with the large BIC iMEP did not change the results (ρ = 0.81, p = 0.0005). **B) In FDI**, CST projection strength did not relate to muscle strength (p = 0.198) but CReST projections did, with larger iMEPs associated with higher MVFs (p < 0.001).

In the distal segment, CST projection strength did not relate to FDI strength (ρ = 0.35, p = 0.198; Fig. 7B). However, CReST projection strength significantly related to FDI strength, with larger iMEPs associated with higher MVFs (ρ = 0. 79, p < 0.001; Fig. 7B).

#### Motor control

In the proximal segment, CST projection strength significantly related to BIC motor control, with larger cMEPs associated with higher motor control indices (ρ = 0.61, p = 0.016; Fig. 8A). Moreover, CReST projection strength showed a strong trend for relating to BIC motor control (ρ = 0. 49, p = 0.063; Fig. 8A).

**Figure 8.**
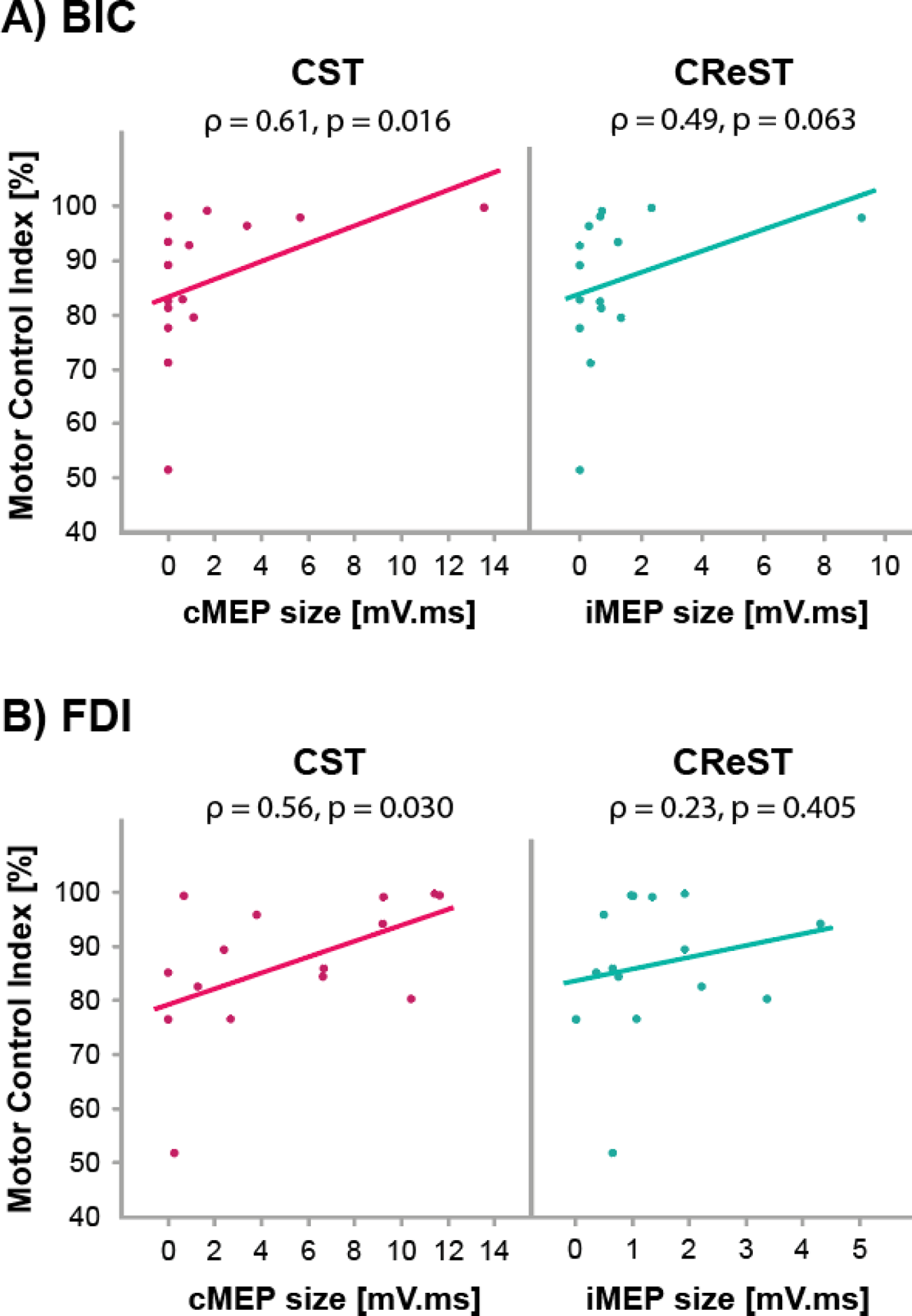
Ipsilesional CST projections strongly relate to paretic motor control. **A) In BIC,** CST projection strength significantly related to BIC motor control, with larger cMEPs associated with higher motor control indices (p = 0.016). CReST projection strength showed a strong trend (p = 0.063) for this relationship. Removal of the subject with the large BIC iMEP reduced this association (ρ = 0.42, p = 0.135). **B) In FDI,** CST projection strength significantly related to FDI motor control, with larger cMEPs associated with higher motor control indices (p = 0.030). CReST projection strength did not relate to FDI motor control (p = 0.405).

In the distal segment, CST projection strength significantly related to FDI motor control, with larger cMEPs associated with higher motor control indices (ρ = 0. 56, p = 0.030; Fig. 8B). CReST projection strength did not relate to FDI control (ρ = 0.23, p = 0.405; Fig. 8B).

#### Muscle individuation

In the proximal segment, CST projection strength significantly related to BIC individuation, with larger cMEPs associated with higher individuation indices (ρ = 0.68, p = 0.005; Fig. 9A). CReST projection strength also significantly related to BIC individuation, with larger iMEPs associated with higher individuation indices (ρ = 0.63, p = 0.011; Fig. 9A)

**Figure 9.**
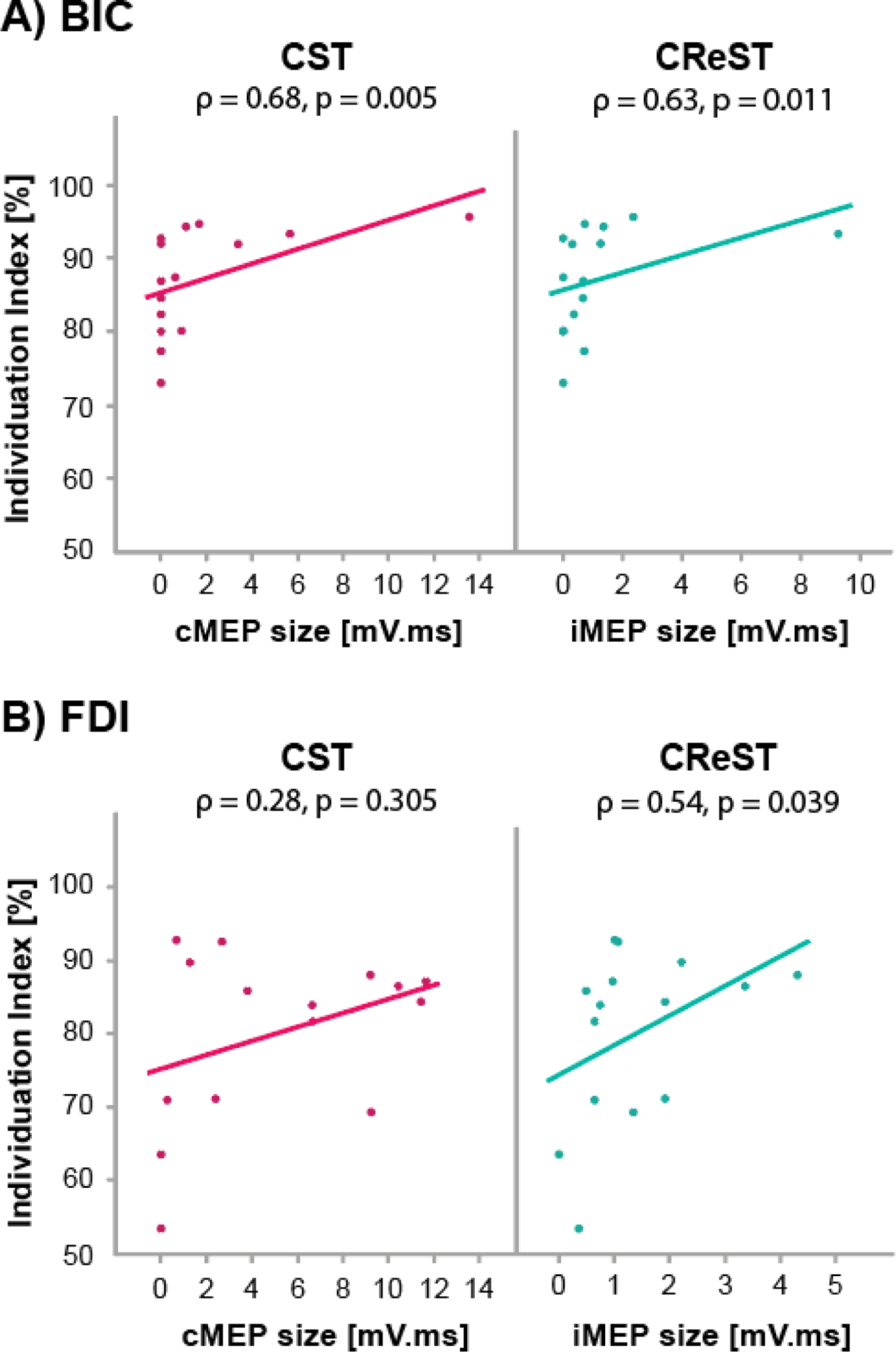
Contralesional CReST projections strongly relate to paretic muscle individuation. **A) In BIC,** CST and CReST projection strengths significantly related to individuation. Larger cMEPs were associated with higher individuation indices (p = 0.005) and larger iMEPs were associated with higher individuation indices (p = 0.011). Removal of the subject with the large BIC iMEP did not change the results (ρ = 0.58, p = 0.031). **B) In FDI,** CST projection strength did not relate to FDI individuation (p = 0.305), but CReST projections did, with larger iMEPs associated with higher individuation indices (p = 0.039).

In the distal segment, CST projection strength did not relate to FDI individuation (ρ = 0.33, p = 0.305; Fig. 9B). However, CReST projection strength was related to FDI individuation, with larger iMEPs associated with higher individuation indices (ρ = 0.54, p = 0.039; Fig. 9B).

Collectively, these results suggest that, for both paretic UE segments, the ipsilesional CST may be a primary contributor to motor control and the contralesional CReST may be a primary contributor to muscle strength and individuation.

#### Functional connectivity and segmental behavior

We explored the relevance of functional pathway connectivity to motor behavior in stroke, but the imbalanced group sizes considerably reduced our confidence in the findings. Assessments are reported in Table 4 for historical comparison.

**Table 4.** Functional pathway connectivity and motor behaviors in stroke subjects. Median (IQR) behaviors are shown for ipsilesional CST and contralesional CReST. Behaviors are split by cMEP or iMEP presence or absence. We note that imbalanced groups limit powered inference about behavioral differences associated with pathway connectivity, although significant differences within muscle and pathway are noted for historical reference. * p < 0.05, median (IQR) shown.

| BIC |  |  |  |  | FDI |  |  |  |
| --- | --- | --- | --- | --- | --- | --- | --- | --- |
|  | Ipsilesional CST |  | Contralesional CReST |  | Ipsilesional CST |  | Contralesional CReST |  |
|  | cMEP+ | cMEP- | iMEP+ | iMEP- | cMEP+ | cMEP- | iMEP+ | iMEP- |
| Subject n | 7 | 8 | 10 | 5 | 13 | 2 | 14 | 1 |
| MVF (N) | 65.7 | 42.9 | 64.7* | 31.0 | 8.6* | 2.5 | 8.4 | 2.6 |
|  | (57.4) | (15.5) | (36.0) | (22.2) | (5.5) | (0.1) | (6.3) | (0) |
| Motor control index (%) | 95.6 | 81.3 | 94.6 | 83.0 | 89.4 | 80.9 | 87.7 | 76.6 |
|  | (16.3) | (19.6) | (18.8) | (26.2) | (17.8) | (8.7) | (17.2) | (0) |
| Individuation index (%) | 93.3* | 83.5 | 92.0 | 80.1 | 85.9* | 58.5 | 85.2 | 63.6 |
|  | (7.3) | (12.7) | (10.4) | (13.5) | (12.5) | (10.1) | (17.4) | (0) |

### Segmental behaviors and descending motor neurophysiology in healthy subjects

For normative comparison, we also examined healthy subjects’ segmental behaviors and their relationships to descending motor pathways. Like stroke subjects, motor control did not significantly differ between the BIC and FDI (z = 1.5, p = 0.145). Like stroke subjects, muscle individuation was worse with FDI than BIC (z = 4.5, p < 0.0001). In terms of pathway-behavior relationships, motor behaviors were mostly unrelated to CST or CReST projection strengths in the BIC (ρ < 0.28, p > 0.146) or FDI (ρ < 0.24 p > 0.226), with a couple of exceptions. For the CReST, larger iMEPs related to better individuation in the BIC (ρ = 0. 47, p = 0.003) and showed a trend for relating to poorer motor control in the FDI (ρ = −0.4, p = 0.062).

## Discussion

In this cross-sectional chronic stroke study, we examined how two major descending motor pathways—the ipsilesional corticospinal tract and the contralesional corticoreticulospinal tract—relate to segmental motor behaviors in the paretic UE. We used quantitative testing to measure strength, motor control, and muscle individuation in a representative muscle of the proximal and distal UE segment. We used TMS to probe the projection strengths of the ipsilesional CST and contralesional CReST to the muscles. We found that, for both UE segments, CST projection strength was critical for motor control and CReST projection strength was critical for muscle strength and individuation. These pathways also shared motor behaviors in the proximal segment, with CST relating to proximal strength and individuation and CReST trending a relationship to motor control. We also found that motor control capabilities were similar for both UE segments, but muscle individuation was poorer for the distal segment. These observations suggest dissociable but collaborative contributions of the CST and CReST to chronic motor behaviors, which vary not only by UE segment but also by their effectiveness.

### Neural Pathways and their relationship to UE deficits in stroke

#### Pathway relationships to segmental strength

In this study, we found that the strength of projections from both the ipsilesional CST and contralesional CReST related to proximal muscle strength. In contrast, a previous chronic stroke study that examined TMS neurophysiology and proximal motor behaviors found no relationship between paretic pectoralis strength and a relative metric of CST versus CReST projection strengths (Schwerin *et al*., 2008). This contradictory finding could be due to the use of the relative pathway metric, which cannot dissociate independent pathway relationships to a behavior. Relative metrics can overlook situations where both pathways similarly relate to a behavior, as we found for CST and CReST, because it detects no relative difference.

We also found that ipsilesional CST projections do not relate to distal strength, in agreement with a previous study that also found no CST relationship to FDI strength (Brouwer & Schryburt-Brown, 2006). Others found positive relationships to grip strength (Pennisi *et al*., 2002; Thickbroom *et al*., 2002, 2004) or composite UE strength (Homberg *et al*., 1991; Cakar *et al*., 2016), but these strength metrics also capture forearm and arm contributions. It is possible that these findings therefore reflect a positive CST association with proximal strength samples, as we found.

We newly observed that stronger CReST projections related to greater paretic FDI strength. Two previous TMS studies found that CReST connectivity (iMEP+) was more prevalent in chronic stroke patients with hand and forearm weakness (Netz *et al*., 1997; Werhahn *et al*., 2003), but neither examined CReST projection strengths or their relationships to individual muscle strength. Although these results have been interpreted by some as a maladaptive influence of the CReST (Netz *et al*., 1997), our findings suggest a benefit.

#### Pathway relationships to motor control

We also examined paretic motor control, finding that stronger ipsilesional CST projections relate to better proximal motor control. These findings may align with a chronic stroke study that examined TMS-evoked responses from both hemispheres and proximal motor control (Hammerbeck *et al*., 2019). Better reaching accuracy was observed in subjects with ipsilesional CST connectivity present (cMEP+) compared to subjects with connectivity absent (cMEP-) but with contralesional responses present. We note that the classification of a contralesional response included both iMEPs and ipsilateral silent periods, which represent distinct neural pathways. These findings therefore could represent better proximal control in the presence of CST connectivity, in the absence of CReST connectivity, and/or in the presence of weak transcallosal inhibition. Our results indicate that the strength of ipsilesional CST projections is key for proximal motor control, and suggest potential additional contributions from CReST.

We also found that stronger ipsilesional CST projections relate to better distal motor control, in agreement with a chronic stroke study that found stronger ipsilesional CST projections relating to faster finger tapping (Cakar *et al*., 2016). We found no significant relationship between CReST projections and distal motor control. To our knowledge, this relationship has not been previously examined in chronic stroke.

#### Pathway relationships to muscle individuation

We also examined paretic muscle individuation, or the degree to which non-instructed muscles are inactive during task performance. We found that stronger ipsilesional CST and contralesional CReST projections relate to better proximal individuation. These findings may align with the TMS study discussed above (Schwerin *et al*., 2008), which measured non-instructed elbow movement during pectoralis activation with respect to relative pathway projection strengths. This study found poorer pectoralis individuation (i.e., more elbow movement) related to a lower relative metric of CST versus CReST projections, which was interpreted as a stronger CReST relating to worse individuation. However, as noted above, relative metrics cannot dissociate independent pathway-behavior relationships. An alternative interpretation could be that critically weaker CST projections relate to worse individuation, which we observed. To our knowledge, our distal segment findings, showing stronger CReST but not CST projections relating to better FDI individuation, have not been previously examined in chronic stroke.

Our findings of a beneficial relationship between CReST projections and individuation are in contrast to those obtained with an alternate paradigm, weighted shoulder abduction. This maneuver elicits non-instructed flexor activity (i.e. poorer individuation) that is more marked in chronic stroke than healthy subjects (Beer *et al*., 1999; Miller & Dewald, 2012; Lan *et al*., 2017; Wilkins *et al*., 2020). Weighted abduction was found to be associated with increased contralesional EEG activity and non-instructed muscle activation in chronic stroke (McPherson *et al*., 2018; Wilkins *et al*., 2020; Tian *et al*., 2021), which was interpreted as a maladaptive relationship between CReST and flexor synergies. However, EEG cannot distinguish between intracortical, transcallosal, or CReST sources of cortical activity, and these circuits are known to be hyperexcitable in chronic stroke (Schambra *et al*., 2015; Taga *et al*., 2021; Hayes *et al*., 2023). Probing these distinct pathways directly with TMS during shoulder weighting could provide disambiguation.

#### Motor behaviors in proximal and distal UE segments

We also investigated whether there are segmental differences in behavioral motor deficits, comparing motor control and muscle individuation between the BIC and FDI. An important consideration was that the assessments would appraise the segments equally and not introduce task-related differences. We therefore used identical testing paradigms, matching task parameters, comparable muscle activation, and identical outcome measures to make unbiased comparisons.

We found similar motor control capabilities by the BIC and FDI, suggesting that single-joint motor control deficits have no particular segmental bias in chronic stroke. Comparison with previous stroke studies is challenging as most examined only one segment (Levin, 1996; Cirstea *et al*., 2003; Hermsdörfer *et al*., 2003; Wolbrecht *et al*., 2018), or used different behavioral tasks and outcome measures to compare control in each segment. For example, investigators used proximal reaching and distal grasping to examine chronic motor control, finding worse distal outcomes (Wenzelburger *et al*., 2005; Lang *et al*., 2006). Although these movements are functionally relevant to each segment, they require different testing parameters (e.g. targets, movement ranges, movement speeds, and error tolerance) that alter performance demands and may influence segmental read-outs.

We also found that muscle individuation was poorer during FDI task performance than BIC performance; in other words, non-instructed muscles throughout the UE showed greater activity during controlled movement at the index finger than elbow. These results differ from other chronic stroke studies that found worse individuation with proximal movement (Zackowski *et al*., 2004; McPherson & Dewald, 2022) or no segmental difference (Lang & Beebe, 2007). This disagreement may reflect differences in how non-instructed muscle activity was elicited. Our skilled motor task required submaximal activation of the instructed muscle and active range of motion (AROM); others used maximal activation and full AROM. Given that greater instructed muscle activation provokes greater non-instructed muscle activation (Xu *et al*., 2017; McPherson *et al*., 2018; Tian *et al*., 2021), subtler individuation differences may be lost to saturation by more intense task demands. Alternatively, isolated FDI movement may have elicited an active stabilization strategy, despite restraints, to minimize interaction torques in non-instructed joints (Gribble & Ostry, 1999). If this were the case, however, we would have expected predominant flexor activation with the FCR and FDS to stabilize the wrist and hand, not the extensor predominance that we observed.

#### Proposed anatomical-functional framework

We found that the ipsilesional CST generally related to paretic UE motor control and the contralesional CReST generally related to paretic UE strength and individuation. Each pathway also had complementary relationships to motor behaviors in the proximal segment (Fig. 10). How do these pathway-behavior observations relate to the known neuroanatomical features of the CST and CReST?

**Figure 10.**
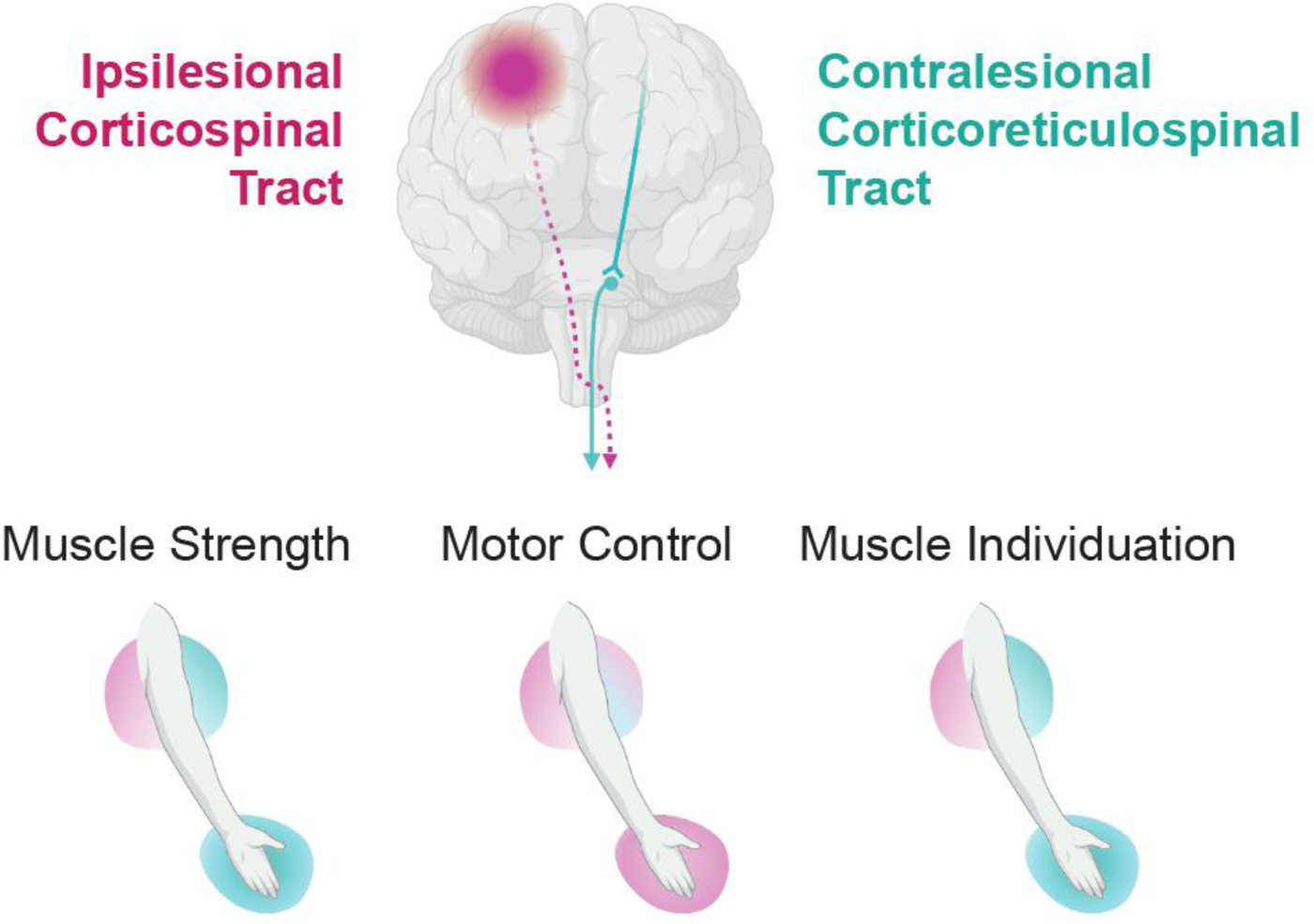
Proposed CST and CReST mediation of chronic motor behaviors. After the CST is injured in stroke (purple dashed line), the contralesional CReST (blue line) may take on a role of mediating motor behavior of the UE. This schematic shows CST and CReST projection strengths related to segmental motor behaviors, with the strength of associations scaled with color. We found an association between stronger ipsilesional CST projection and better motor control in both UE segments, while we found an association between stronger contralesional CReST projections and better strength and individuation in both UE segments. Importantly, we found that both pathways have shared associations with behaviors in the proximal UE segment: Higher projection strength of both pathways were associated with better strength and individuation in the proximal UE segment. (CST, corticospinal tract; CReST, corticoreticulospinal tract; BIC, biceps; FDI, first dorsal osseous). Created with BioRender.com.

A main theme arising from anatomical tracing, electrophysiological, and neurochemical studies in animals is that the pathways have much in common. First, both pathways originate from shared primary and secondary motor areas (He *et al*., 1993, 1995; Matsuyama & Drew, 1997; Boudrias *et al*., 2010a; Boudrias *et al*., 2010b; Fisher *et al*., 2012; Fregosi *et al*., 2017), pointing to their shared capacity to generate voluntary movement. Second, both pathways collateralize rostrocaudally to the upper and lower cervical spinal cord, and both diverge within the cord to terminate on medial and lateral motoneuronal pools (Kuypers *et al*., 1962; Peterson *et al*., 1975; Shinoda *et al*., 1979; Matsuyama *et al*., 1997; McKiernan *et al*., 1998; Riddle *et al*., 2009; Sinopoulou *et al*., 2022; Morecraft *et al*., 2023). This extensive branching points to their shared capacity to affect proximal and distal UE muscles and agonist-antagonist pairs. Third, both pathways terminate most densely in the spinal intermediate laminae (Kuypers *et al*., 1962; Kuypers & Brinkman, 1970; Dum & Strick, 1996; Morecraft *et al*., 2013; Morecraft *et al*., 2023). The intermediate laminae contain dendrites of ventral horn motoneurons and many types of excitatory and inhibitory interneurons: local interneurons with ipsilateral projections, commissural interneurons with contralateral projections, and long and short propriospinal neurons with bilateral projections to multiple spinal segments (Lawrence *et al*., 1985; Zholudeva *et al*., 2021; Sinopoulou *et al*., 2022). This elaborate connectivity points to the shared capacity of the pathways to influence numerous muscles throughout the UE. In sum, both the CST and CReST connect to multiple spinal levels and distributed motoneuron pools, and both pathways can exert excitatory and inhibitory influences via interneuronal targets. These pronounced similarities indicate the potential for redundancy and partnership between pathways.

There are, however, some unique pathway features that suggest preferential management of particular motor behaviors, in alignment with our behavioral observations. We found an association between ipsilesional CST projections and UE motor control. Motor control requires the precise and rapid activation of task-specific motor units, believed to be facilitated by corticomotoneuronal (CM) projections within the CST (McKiernan *et al*., 1998; Lemon & Griffiths, 2005). In macaques, this fast-conducting path terminates on and facilitates spinal motoneurons serving UE muscles, particularly distal ones (Porter, 1985; McKiernan *et al*., 1998; Rathelot & Strick, 2009; Morecraft *et al*., 2013). However, despite this distal predilection, we did not observe poorer motor control in the proximal segment. Interestingly, we did observe a strong trend between contralesional CReST projections and proximal motor control. In macaques, fast-conducting reticulospinal projections facilitate motoneurons serving UE muscles, particularly proximal ones (Davidson *et al*., 2007; Riddle *et al*., 2009; Soteropoulos *et al*., 2012; Hirschauer & Buford, 2015), and reticular neurons modulate their firing during proximal UE movement preparation (Buford & Davidson, 2004). It is possible that proximally strong CReST projections supplement proximally weaker CM projections, evening out motor control ability for both segments.

Separately, we found an association between contralesional CReST projections and UE muscle strength and individuation. Sustained force production necessitates the coordinated activation of agonists and suppression of antagonists, while individuating movement requires widespread suppression of non-instructed muscles. Although the CST and CReST both have access to these distributed muscle groups, studies in rats and macaques has found that their capacity for excitation and inhibition differ. Nearly all CST terminals are excitatory (Du Beau *et al*., 2012) and the majority connect to excitatory spinal interneurons (Sinopoulou *et al*., 2022). The majority of reticulospinal terminals are also excitatory, but a sizable minority of terminals are directly inhibitory (Holstege, 1991; Du Beau *et al*., 2012). Moreover, in cats and macaques, reticulospinal activation elicits reciprocal excitation and suppression of spinal motoneurons and their UE muscles (Jankowska *et al*., 1968; Peterson *et al*., 1979; Davidson & Buford, 2006; Schepens & Drew, 2006; Davidson *et al*., 2007). It is thus conceivable that the CReST, with its inhibitory capacity, could mediate the muscle suppression needed to support strength and individuation.

We also found that individuation was worse with distal movements and better with proximal movements. Although this could be due to the reticulospinal predilection for proximal muscle motoneurons (Davidson *et al*., 2007; Riddle *et al*., 2009; Soteropoulos *et al*., 2012; Hirschauer & Buford, 2015), we also found that stronger ipsilesional CST projections related to better proximal individuation. In rats and macaques, the evolutionarily older CST projections arising from rostral M1 and secondary motor areas preferentially terminate in the upper cervical cord, mediating proximal UE muscles (He *et al*., 1993, 1995; Morecraft *et al*., 2019; Sinopoulou *et al*., 2022; Morecraft *et al*., 2023). It is possible that this proximally stronger CST projection could supplement CReST projections, providing additional support for individuation during proximal movement.

How the CST and CReST interact to manage motor behaviors after stroke is not fully clear. In rat, cat, and macaque, up to 30% of CST neurons send collaterals to the pontomedullary reticular formation as they descend to the spinal cord (Brodal, 1981; Keizer & Kuypers, 1984; Keizer & Kuypers, 1989; Sinopoulou *et al*., 2022). Moreover, the CST and CReST share convergent inputs on interneurons in the spinal cord (Riddle & Baker, 2010). These observations suggest anatomic substrates that have capacity to communicate and integrate descending signals.

### Limitations

Our study has some limitations to consider. Because of our behavioral testing criteria, our results reflect pathway-behavior relationships in mild-to-moderately impaired stroke subjects. It is possible that greater stroke damage to CST could drive mediation of behaviors more fully to the contralesional CReST (Di Pino *et al*., 2014). Future studies could examine a broader range of impairment, although behavioral testing in subjects with severe paresis can be challenging. In addition, we used TMS to examine pathways during rest or submaximal isometric force production, not during performance of the motor behaviors. This approach controlled for the degree of muscle pre-activation, which can affect measurements of pathway strength. Thus, our projection strength read-outs indicate the functional capacity of the pathway to support a behavior, but may not reflect how the pathway is used during the behavior. Future studies could examine CST and CReST projection strengths during behavioral performance, although the level of muscle activity requires attention to ensure controlled pathway samples. We also examined two muscles—the BIC and FDI—in the proximal and distal segment. These muscles may not be representative of all segmental muscles and pathway roles may vary by actuator function, even within segment (Davidson *et al*., 2007; Baker, 2011). Testing a broader array of UE muscles, including flexor-extensor and agonist-antagonist pairs, would provide a comprehensive characterization of pathway-behavior relationships. Finally, and critically, our findings were based on chronic observations, and therefore do not reveal pathway roles in motor recovery, which occurred months to years prior. The longitudinal observation of neurophysiological and behavioral changes in subacute stroke is required to evaluate this recovery relationship.

## Conclusions

In stroke subjects with chronic motor deficits, we found that the projection strengths of two key motor pathways—the ipsilesional CST and contralesional CReST—have unique and shared relationships to paretic UE motor behaviors. Stronger ipsilesional CST projections were linked to superior motor control in both UE segments, whereas stronger contralesional CReST projections were linked to superior muscle strength and individuation in both UE segments. Notably, both pathways also shared associations with behaviors in the proximal segment. These results suggest that each pathway has specialized contributions to chronic motor behaviors but also work together, providing human evidence for established theories (Keizer & Kuypers, 1984; Baker, 2011). This dual-management descending motor system may be ethologically advantageous to support chronic motor function after stroke.

## Acknowledgements

The authors thank our healthy controls and chronic stroke participants for their assistance during data collections.

